# AtDREB2G is a novel regulator of riboflavin biosynthesis under low-temperature stress and abscisic acid treatment in *Arabidopsis thaliana*

**DOI:** 10.1101/2023.09.06.556598

**Authors:** Junya Namba, Miho Harada, Yuina Toda, Takanori Maruta, Takahiro Ishikawa, Shigeru Shigeoka, Kazuya Yoshimura, Takahisa Ogawa

**Affiliations:** Department of Life Science and Biotechnology, Faculty of Life and Environmental Sciences, Shimane University, 1060 Nishikawatsu, Matsue, Shimane 690-8504, Japan; Graduate School of Natural Science and Technology, Shimane University, 1060 Nishikawatsu, Matsue, Shimane 690-8504, Japan; Department of Advanced Bioscience, Faculty of Agriculture, Kindai University, Nakamachi, Nara 631-8505, Japan; Institute of Agricultural and Life Sciences, Academic Assembly, Shimane University, 1060 Nishikawatsu, Matsue, Shimane 690-8504, Japan; Experimental Farm, Kindai University, Yuasa, Wakayama, 643-0004, Japan; Department of Food and Nutritional Science, College of Bioscience and Biotechnology, Chubu University, 1200 Matsumoto-cho, Kasugai, Aichi, 487-8501, Japan

**Keywords:** riboflavin biosynthesis, cofactor, regulatory factor, DREB, environmental stress, abscisic acid

## Abstract

Riboflavin (RF) serves as a precursor to FMN and FAD, crucial cofactors in various metabolic processes. Strict regulation of cellular flavin homeostasis is imperative, yet information regarding the factors governing this regulation remains largely elusive. In this study, we first examined the impact of external flavin treatment on the *Arabidopsis* transcriptome to identify novel regulators of cellular flavin levels. Our analysis revealed alterations in the expression of 49 putative transcription factors. Subsequent reverse genetic screening highlighted a member of the Dehydration-Responsive Element Binding (DREB) family, AtDREB2G, as a potential regulator of cellular flavin levels. Knockout mutants of *AtDREB2G* (*dreb2g*) exhibited reduced flavin levels and decreased expression of RF biosynthetic genes compared to wild-type plants. Conversely, conditional overexpression of *AtDREB2G* led to an increase in the expression of RF biosynthetic genes and elevated flavin levels. In wild-type plants, exposure to low temperatures and abscisic acid treatment stimulated enhanced flavin levels and upregulated the expression of RF biosynthetic genes, concomitant with the induction of *AtDREB2G*. Notably, these responses were significantly attenuated in *dreb2g* mutants. Our findings establish AtDREB2G as a novel positive regulator of flavin biosynthesis in *Arabidopsis*, particularly under conditions of low temperature and abscisic acid treatment.

## Introduction

Riboflavin (RF), commonly recognized as vitamin B_2_, is a water-soluble vitamin primarily found *in vivo* in the form of derivatives, namely flavin adenine dinucleotide (FAD) and flavin mononucleotide (FMN). FAD and FMN function as crucial redox cofactors across all living organisms, playing essential roles in a diverse array of metabolic processes. These processes encompass mitochondrial electron transport, the antioxidant network, protein folding, and chromatin remodeling, as well as the metabolisms of nucleotides, amino acids, other cofactors, and a multitude of biologically significant compounds (Lynch et al., 2018). Additionally, they are indispensable for specialized functions such as blue light signaling and photosynthesis in plants (Briggs and Olney, 2001; Roeber et al., 2021; Eggers et al., 2021; Lynch et al., 2022). Consequently, the regulation of cellular flavin levels is of paramount importance.

Riboflavin is synthesized *de novo* in plants, bacteria, and yeast, but not in animals. In bacteria and plants, its biosynthesis follows a common pathway, commencing with one molecule of GTP and two of ribulose 5-phosphate as initial precursors (Hasnain et al., 2013; Sa et al., 2016). Initially, GTP is converted to 5-amino-6-ribitylamino-2,4 (1*H*,3*H*) pyrimidinedione (ARP) through a series of four enzymatic reactions catalyzed by GTP cyclohydrolase II (AtRibA1; At5g64300 in *Arabidopsis thaliana*), 2,5-diamino-6-ribosylamino-4-(3H)-pyrimidinone 5’-phosphate deaminase (PyrD; At4g20960), 5-amino-6-ribosylamino-2,4(1H,3H)-pyrimidinedione 5’-phosphate reductase (PyrR; At3g47390), and 5-amino-6-ribitylamino-2,4(1H,3H) pyrimidinedione 5’-phosphate phosphatase (AtPyrP2; At4g11570) (Herz et al., 2000; Hedtke et al., 2012; Hasnain et al., 2013; Sa et al., 2016). In parallel, ribulose 5-phosphate is converted into 3,4-dihydroxy-2-butanone 4-phosphate by 3,4-dihydroxy-2-butanone-4-phosphate synthase. In *Arabidopsis*, this reaction is catalyzed by a bifunctional enzyme encoded by *AtRibA1*, which exhibits both GTP cyclohydrolase II and 3,4-dihydroxy-2-butanone-4-phosphate synthase activities (Herz et al., 2000; Hedtke et al., 2012). These two intermediates subsequently undergo condensation to produce 6,7-dimethyl-8-ribityllumazine in a reaction catalyzed by lumazine synthase (LS; At2g44050) (Jordan et al., 1999; Xiao et al., 2004). The final step in the RF biosynthetic pathway is mediated by RF synthase (RS; At2g20690) (Fischer et al., 2005). FMN is generated from RF through ATP-dependent phosphorylation by RF kinases, and FAD is subsequently formed from FMN *via* ATP-dependent adenylation by FAD synthetase (Roje, 2007; Lynch et al., 2018).

Based on bioinformatic and experimental evidence, it is likely that RF biosynthesis occurs exclusively within plastids in plants (Roje, 2007; Gerdes et al., 2012; Eggers et al., 2021). However, the production of FMN and FAD likely takes place not only in plastids but also in the cytosol and mitochondria. The activities of these enzymes have been detected in several plant species, and genes encoding some of these enzymes have been cloned from *Arabidopsis*. Cytosolic RF kinase has been found to possess both RF kinase activity and FMN hydrolase activity (Sandoval and Roje, 2005; Roje, 2007). Additionally, the activities of RF kinase and FAD synthetase, as well as FMN and FAD hydrolase activities, have been observed in plastids (Sandoval et al., 2008). We have demonstrated that an *Arabidopsis* NUDX (nucleotide diphosphate linked to some other moiety X) hydrolase, AtNUDX23, is localized in chloroplasts and exhibits pyrophosphohydrolase activity towards FAD, enabling the hydrolysis of FMN and AMP (Ogawa et al., 2008).

The exogenous application of RF to plants has been documented to trigger resistance mechanisms against pathogens by priming defense responses (Dong and Beer, 2000; Zhang et al., 2009). Conversely, a reduction in intrinsic RF content, achieved through the ectopic expression of turtle riboflavin-binding protein (RfBP), has been associated with increased levels of H_2_O_2_ and enhanced plant resistance to bacterial pathogens (Deng et al., 2011). Suppression of *AtRibA1* expression in *Arabidopsis* antisense lines resulted in a bleached leaf phenotype (Hiltunen et al., 2012). In a previous study, we observed that exogenous treatment of *Arabidopsis* leaves with flavins did not alter the levels of FMN and FAD, despite an enhancement in RF levels, their precursor. Maintaining constant FAD and FMN levels was likely achieved through transcriptional feedback regulation of flavin metabolism (Maruta et al., 2012). These findings underscore the importance of stringent control over cellular flavin levels, as they can influence both physiological and pathological responses. In this study, we aimed to identify transcriptional regulators of flavin metabolism in plants by combining preliminary transcriptome analysis and reverse genetic screening. Our approach successfully pinpointed AtDREB2G, a member of the Dehydration-Responsive Element Binding (DREB) family, as a positive regulator of RF biosynthesis in response to low temperatures and abscisic acid (ABA).

## Results

### Identification of AtDREB2G as a regulator of flavin homeostasis

In our previous work (Maruta et al., 2012), the cellular levels of RF in *Arabidopsis* increased within an hour of exogenous treatment with FMN, FAD, and RF. Furthermore, this treatment was found to impact the transcript levels of genes involved in flavin metabolism, indicating that excess FMN and FAD are promptly converted to RF in plant cells, and that flavin-dependent transcriptional regulation occurs within an hour of flavin treatment. Therefore, in this study, we conducted a transcriptome analysis of *Arabidopsis* leaves following an hour of mock (water) or 0.1 mM FAD treatment using Affymetrix ATH1 GeneChip arrays (GEO accession number GSE169061) as an initial screening. This analysis identified 262 genes with increased expression and 198 genes with decreased expression upon FAD treatment (more than two-fold compared to the mock). Upon classification of these genes based on the broad GO ontology categories developed at TAIR (http://www.geneontology.org/GO.slims.shtml), we identified 49 genes encoding putative transcription factors (Supplemental Table S1). Subsequently, we focused on exploring the functions of these transcription factors.

To ascertain the involvement of these proteins in the regulation of flavin metabolism, we conducted a comprehensive analysis of their influence on flavin levels using knockout mutants. We were able to obtain T-DNA insertion mutant lines for 19 out of the 49 genes from the Arabidopsis Biological Resource Center (https://abrc.osu.edu) (Supplemental Tables 1 and 2). By comparing flavin levels in these knockout mutants to those in wild-type plants, we identified four mutants that exhibited significant alterations in flavin levels (Supplemental Fig. S1). Among these mutants, the *AtDREB2G* mutants, *dreb2g-1* (CS452041), displayed the most pronounced impact on flavin levels. Specifically, the levels of FMN and RF were significantly reduced to 57% and 51% of the values observed in wild-type plants (Figs. 1A and 1B). Although the levels of FAD in the *dreb2g-1* mutants also exhibited a decreasing trend (87%) compared to those in wild-type plants, the difference was not statistically significant. Additionally, transcript levels of three genes involved in RF biosynthesis, namely *AtRibA1*, *PyrD*, and *LS*, were significantly reduced in *dreb2g-1* mutants when compared to wild-type plants (Fig. 1C). We also obtained another T-DNA insertion line, *dreb2g-2* (CS878257), which similarly demonstrated reduced RF and FMN levels in comparison to wild-type plants (Figs. 1A and 1B).

**Fig. 1.**
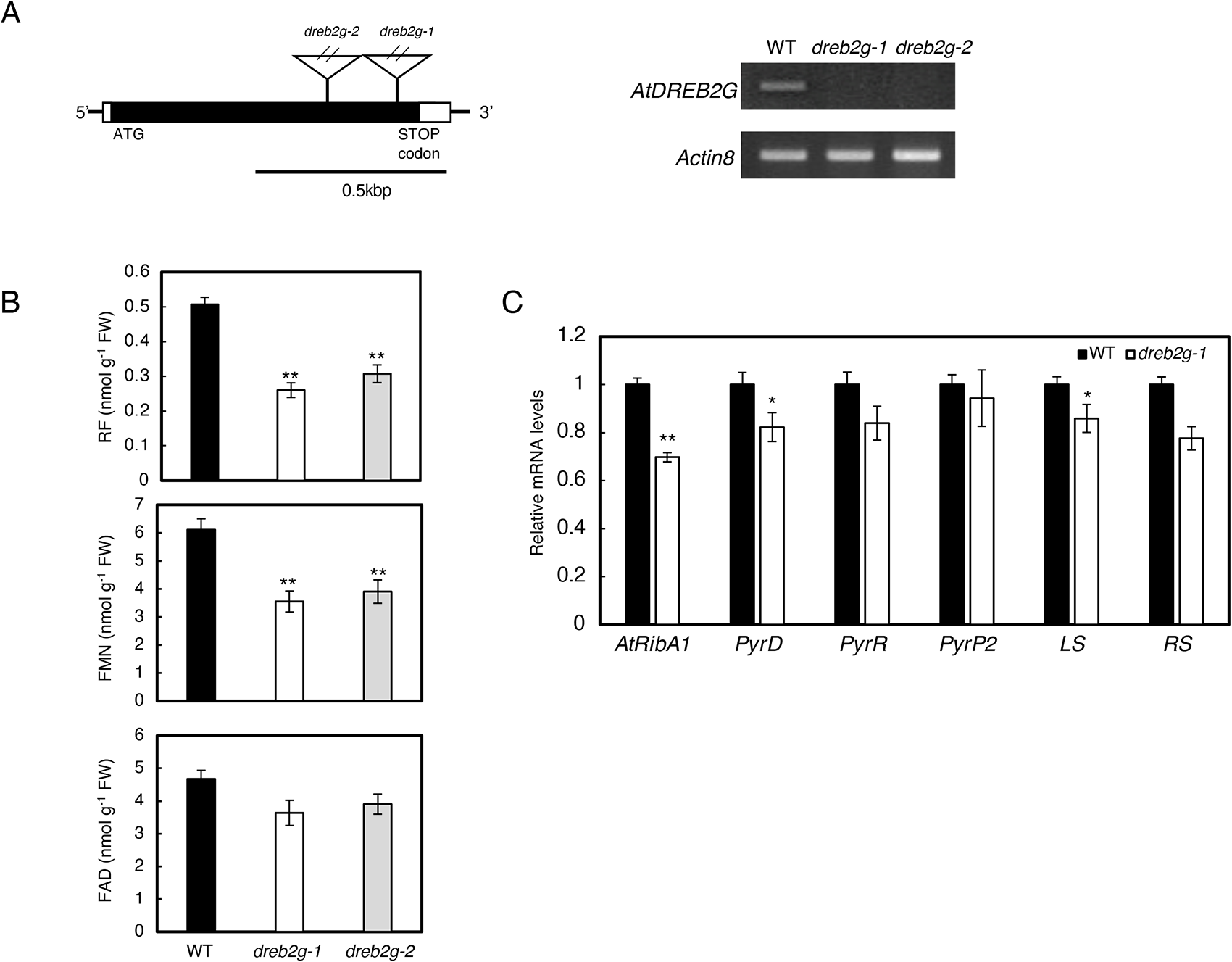
Flavin metabolism in knockout mutants of *AtDREB2G*. Wild-type and *dreb2g* plants were cultivated on 1/2 MS medium supplemented with 1% sucrose for two weeks under normal conditions. Subsequently, leaves were collected four hours after illumination. (A) T-DNA insertion sites, *dreb2g-1* (CS452041) and *dreb2g-2* (CS878257), in the *AtDREB2G* gene and semi-quantitative RT-PCR analysis illustrating the transcription of *AtDREB2G* in both wild-type and mutant plants. T-DNA insertion sites are denoted by triangles, with black and white boxes representing exons and untranslated regions, respectively. (B) Cellular flavin levels and (C) transcript levels of RF biosynthetic genes in the wild-type and mutant plants. The data represent means ± SE from a minimum of three individual experiments (n ≥ 3) involving independently grown plants. Asterisks indicate significant deviations from the values observed in the wild-type plants (Student’s *t*-test, *P < 0.05 and **P < 0.01).

To further substantiate the impact of *AtDREB2G* on flavin metabolism, we utilized an estrogen (ES)-induced conditional expression line (CS2102394), obtained from the ABRC. In this line, the transcription of *AtDREB2G* can be induced by ES treatment. Transcript levels of *AtDREB2G* exhibited a significant increase at 3 and 6 hours after ES treatment, following a time-dependent pattern. Subsequently, they began to decline at the 12-hour mark, ultimately returning to untreated levels by 24 hours (Fig. 2A). Simultaneously with the induction of *AtDREB2G*, we observed a significant upregulation in the transcript levels of *AtRibA1*, *PyrD*, *PyrR*, and *LS* in response to ES treatment (Fig. 2B). Importantly, cellular flavin levels also displayed a significant increase 6 hours after ES treatment (Fig. 2C). These findings unequivocally support the role of *AtDREB2G* as a positive regulator of flavin biosynthesis in *Arabidopsis*.

**Fig. 2.**
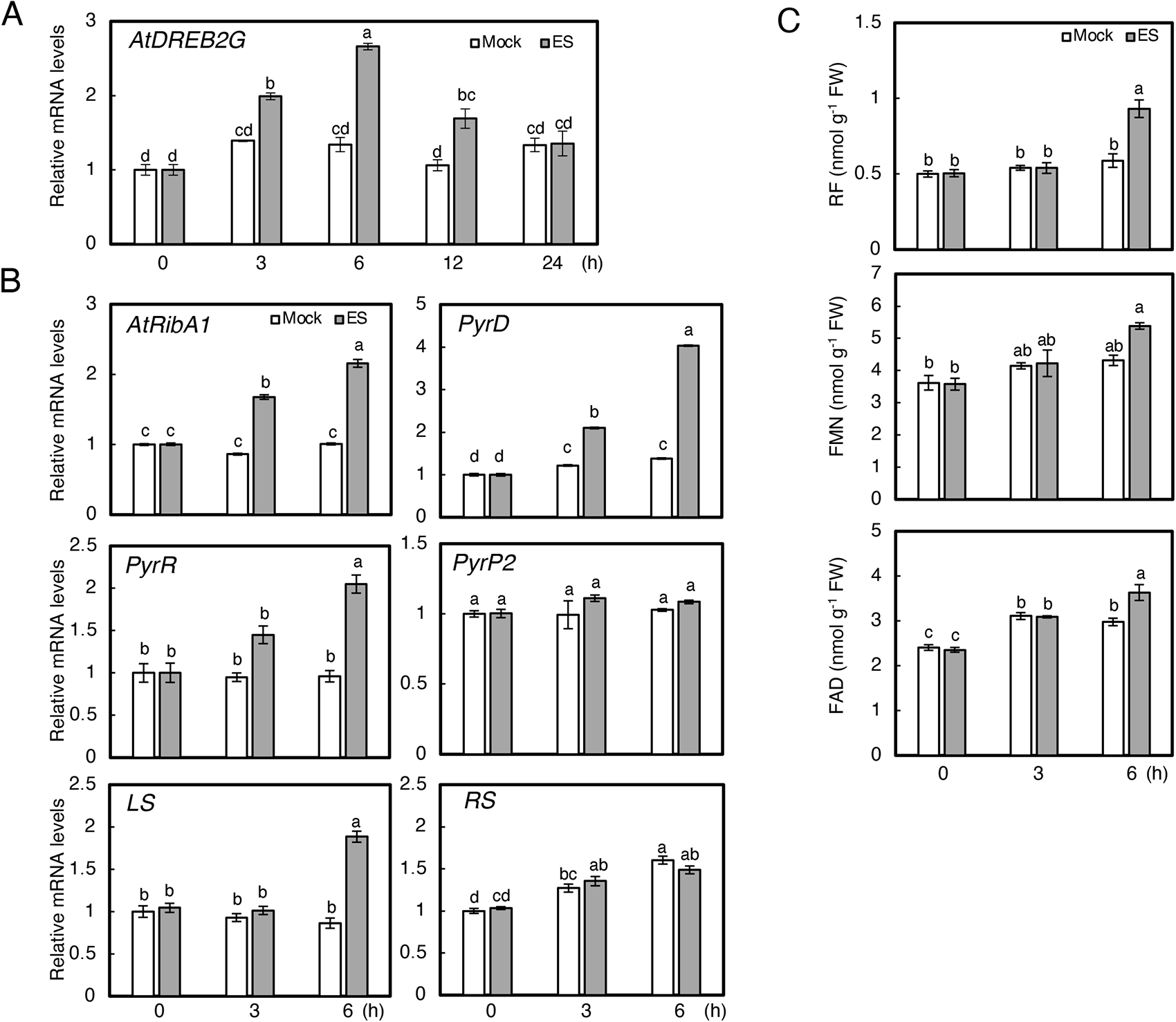
Effects of transient overexpression of *AtDREB2G* on cellular flavin levels and RF biosynthetic gene expressions. Wild-type and transgenic plants, in which *AtDREB2G* expression is induced by estradiol (CS2102394), were cultivated on 1/2 MS medium supplemented with 1% sucrose for two weeks under standard conditions. Subsequently, seedlings were transferred to 1/2 MS medium with or without 2 μM ES. The transcript levels of (A) *AtDREB2G* and (B) RF biosynthetic genes. (C) Cellular flavin levels. The presented data represent means ± SE from a minimum of three individual experiments (n ≥ 3), involving independently grown plants. Different letters indicate significant variations (P < 0.05, Tukey-Kramer test).

### AtDREB2G regulates RF biosynthesis under low-temperature stress and ABA treatment

*AtDREB2G* belongs to the DREB family, a group comprising 15 genes in *Arabidopsis* (Supplemental Fig. S2) (Welling and Palva, 2008). These genes encode plant-specific transcription factors responsible for regulating the expression of stress-inducible genes. They play a crucial role in enhancing resistance to abiotic stresses such as drought, salt, and cold by binding to specific DNA sequences called dehydration-responsive elements (DREs) located in the promoter regions of their target genes (Yamaguchi-Shinozaki and Shinozaki, 1994; Shinozaki and Yamaguchi-Shinozaki, 1997; Liu et al., 1998). AtDREB2G possessed a highly conserved AP2/ERF domain characteristic of the DREB family (Supplemental Fig. S3). However, it lacked a negative regulatory domain (NRD) involved in protein degradation, a feature present in AtDREB2A and GmDREB2A (Sakuma et al., 2006a; Mizoi et al., 2013) (Supplemental Fig. S3). Among the DREB family members, *AtDREB1A*, *1B*, and *1C* genes have been shown to exhibit rapid transcriptional induction in response to low-temperature stress (Shinwari et al., 1998). Conversely, the expression of *AtDREB2A* has been reported to increase in response to drought, salt, and high- and low-temperature stresses (Liu et al., 1998; Sakuma et al., 2006a; 2006b). Furthermore, the expression of *AtDREB2A* and *2E* genes has been demonstrated to be upregulated by ABA treatment, a hormone pivotal in enhancing plant tolerance to drought, salinity, and low-temperature stress (Sakuma et al., 2002).

Although the physiological role of *AtDREB2G* as well as its relationship with cellular flavin levels in abiotic stress responses remains largely uncharted, given the roles of other DREB proteins in stress acclimation, we posited that *AtDREB2G* regulates flavin metabolism under abiotic stress conditions. Thus, we investigated the impact of abiotic stresses (low and high temperatures) and ABA treatments on *AtDREB2G* expression and flavin metabolism. In the subsequent experiments, we assessed the transcript levels of three genes, *AtRibA1*, *PyrD*, and *LS*, pivotal in RF biosynthesis, as their transcription was altered in both *dreb2g* and ES-inducible lines (Figs. 1 and 2). In wild-type plants, the transcription of *AtDREB2G* and RF biosynthetic genes exhibited significant time-dependent induction following a 6-hour low-temperature treatment (4°C), concomitant with a notable rise in cellular flavin levels (Figs. 3A-3C). Conversely, in *dreb2g-1* mutants, the induction of RF biosynthetic genes was notably inhibited compared to that in wild-type plants (Figs. 3A and 3B). The levels of cellular flavins in *dreb2g-1* mutants also remained low even after low-temperature treatment (Figs. 3C).

**Fig. 3.**
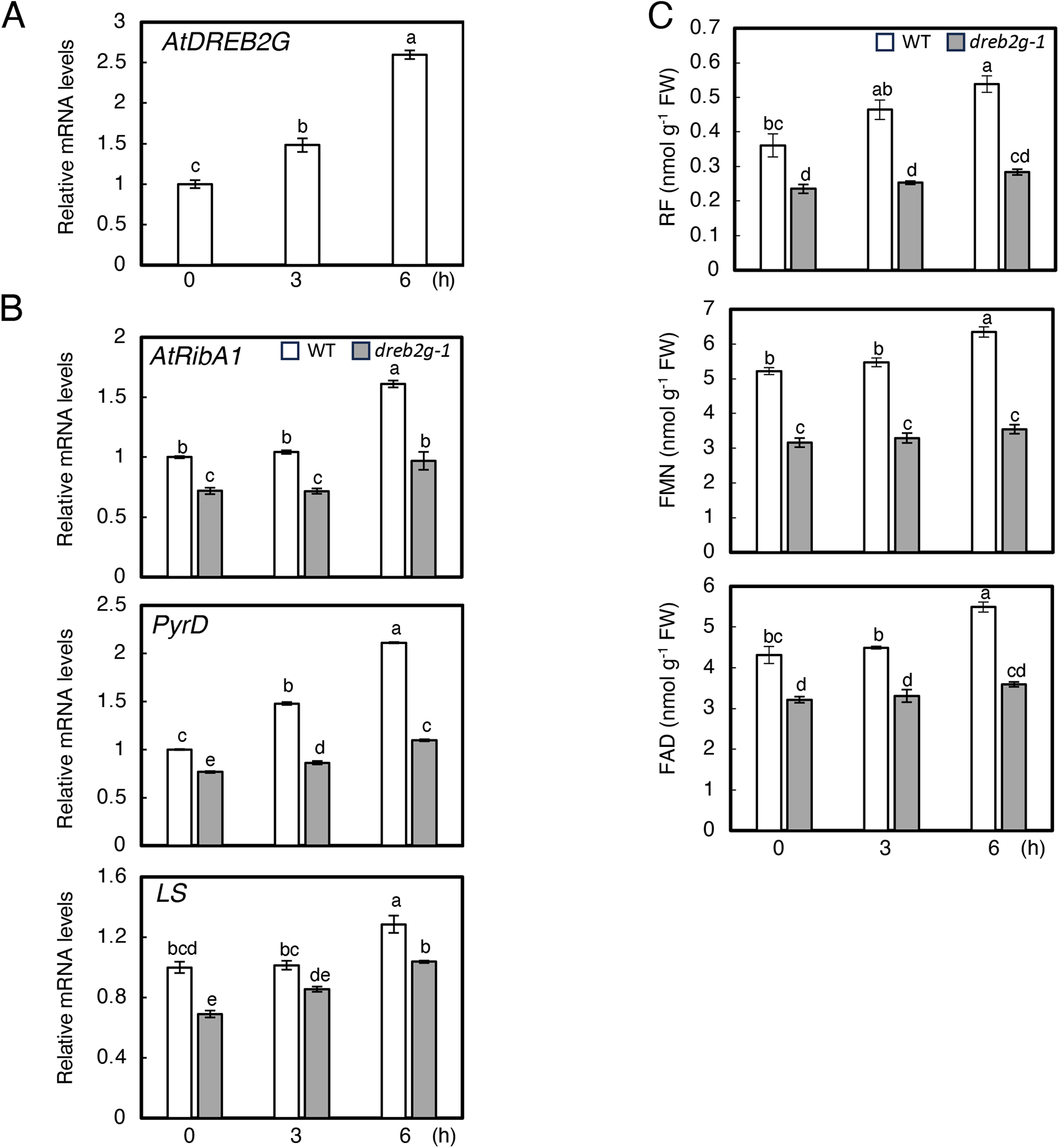
Transcription of *AtDREB2G* and RF biosynthetic genes and cellular flavin levels in the wild type and *dreg2g-1* mutants under low-temperature stress. Wild-type and *dreb2g-1* plants were cultivated on 1/2 MS medium with 1% sucrose for two weeks under standard conditions. Subsequently, seedlings were transferred to 4°C (100 µmol photons/m2/s) for 6 hours. The transcript levels of (A) *AtDREB2G* and (B) RF biosynthetic genes. (C) Cellular flavin levels. The presented data represent means ± SE from a minimum of three individual experiments (n ≥ 3) involving independently grown plants. Different letters indicate significant differences (P < 0.05, Tukey-Kramer test).

The *AtDREB2G* transcription was significantly induced by high-temperature treatment (37°C) in wild-type plants. However, the transcription of *AtRibA1* and *PyrD* remained unaltered in both wild-type and *dreb2g-1* plants (Supplemental Fig. S4). The transcription of *LS* was significantly increased in both wild-type and *dreb2g-1* plants following 6 hours of high-temperature treatment. Cellular flavin levels in both sets of plants tended to decrease after high-temperature treatment, although the differences between wild-type and *dreb2g-1* plants were not statistically significant (Supplemental Fig. S4).

Transcript levels of *AtDREB2G* and RF biosynthetic genes in wild-type plants exhibited significant induction at 24 hours after ABA treatment (50 μM) (Figs. 4A and 4B). Similarly, cellular RF levels in wild-type plants were significantly augmented by ABA treatment (Fig. 4C). Conversely, in *dreb2g-1* mutants, transcript levels of RF biosynthetic genes and RF levels after ABA treatment were notably suppressed compared to those in wild-type plants (Figs. 4B and 4C). Collectively, our findings substantiate that the expression of *AtDREB2G* is transcriptionally induced by low temperature and ABA, thereby governing flavin metabolism.

**Fig. 4.**
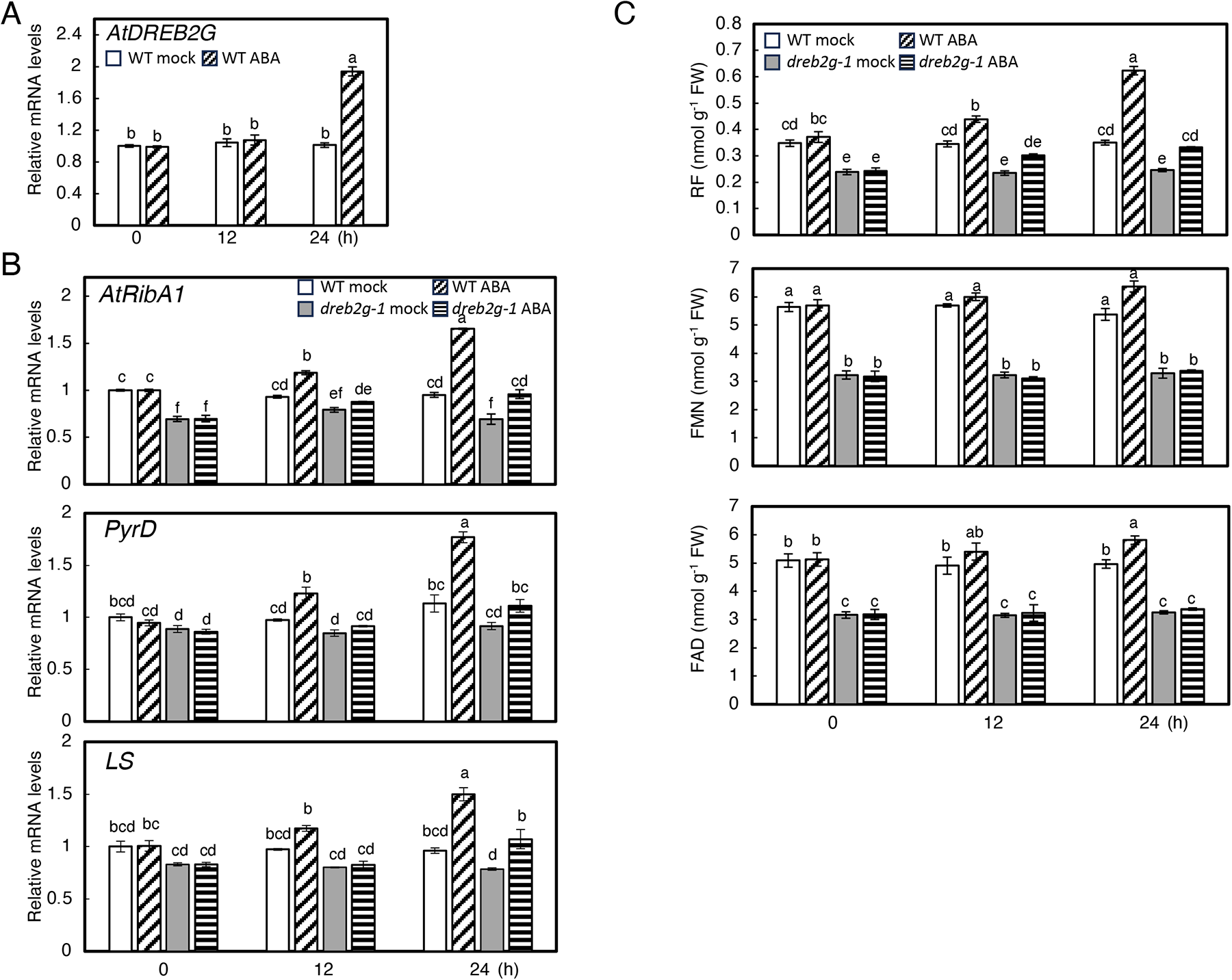
Transcription of *AtDREB2G* and RF biosynthetic genes and cellular flavin levels in the wild type and *dreg2g-1* mutants under ABA treatment. Wild-type and *dreb2g-1* plants were cultivated on 1/2 MS medium with 1% sucrose for two weeks under standard conditions. Subsequently, seedlings were transferred to 1/2 MS medium with or without 50 μM ABA. The transcript levels of (A) *AtDREB2G* and (B) RF biosynthetic genes. (C) Cellular flavin levels. The presented data represent means ± SE from a minimum of three individual experiments (n ≥ 3) involving independently grown plants. Different letters indicate significant differences (P < 0.05, Tukey-Kramer test).

### Stress tolerance and ABA sensitivity of AtDREB2G-overexpressing plants

The induction of *AtDREB2G* by low temperatures led us to hypothesize that this transcription factor plays a pivotal role in stress acclimation. To thoroughly investigate this hypothesis, we generated transgenic *Arabidopsis* plants with constitutive overexpression of *AtDREB2G* in both wild-type (OE-*DREB2G*) and *dreb2g-1* backgrounds (*dreb2g-1*/*DREB2G*) (Figs. 5A and 5B). We also introduced an empty vector into both wild-type plants and *dreb2g-1* mutants as control lines (control and *dreb2g-1*/VC, respectively). The transcript levels of *AtDREB2G* in OE-*DREB2G* and *dreb2g-1*/*DREB2G* lines were increased approximately 200 to 2000-fold compared to those in the control plants (Figs. 5B). Both OE-*DREB2G* and *dreb2g-1*/*DREB2G* plants exhibited robust growth, akin to wild-type and control plants (Fig. 5A). In OE-*DREB2G* plants, the transcript levels of *AtRibA1* and *LS*, as well as cellular RF levels, exhibited slight but significant increases compared to those in the control plants (Figs. 5C and 5D). Furthermore, in *dreb2g-1*/*DREB2G* plants, the transcript levels of genes involved in RF synthesis and the levels of cellular RF and FAD were significantly elevated compared to those in the *dreb2g-1*/VC plants. Consequently, cellular flavin levels, except for FMN, in *dreb2g-1* mutants were restored to the same level as those in the control plants through *AtDREB2G* overexpression (Figs. 5C and 5D). These findings provide additional evidence for the role of this transcription factor in regulating flavin metabolism.

**Fig. 5.**
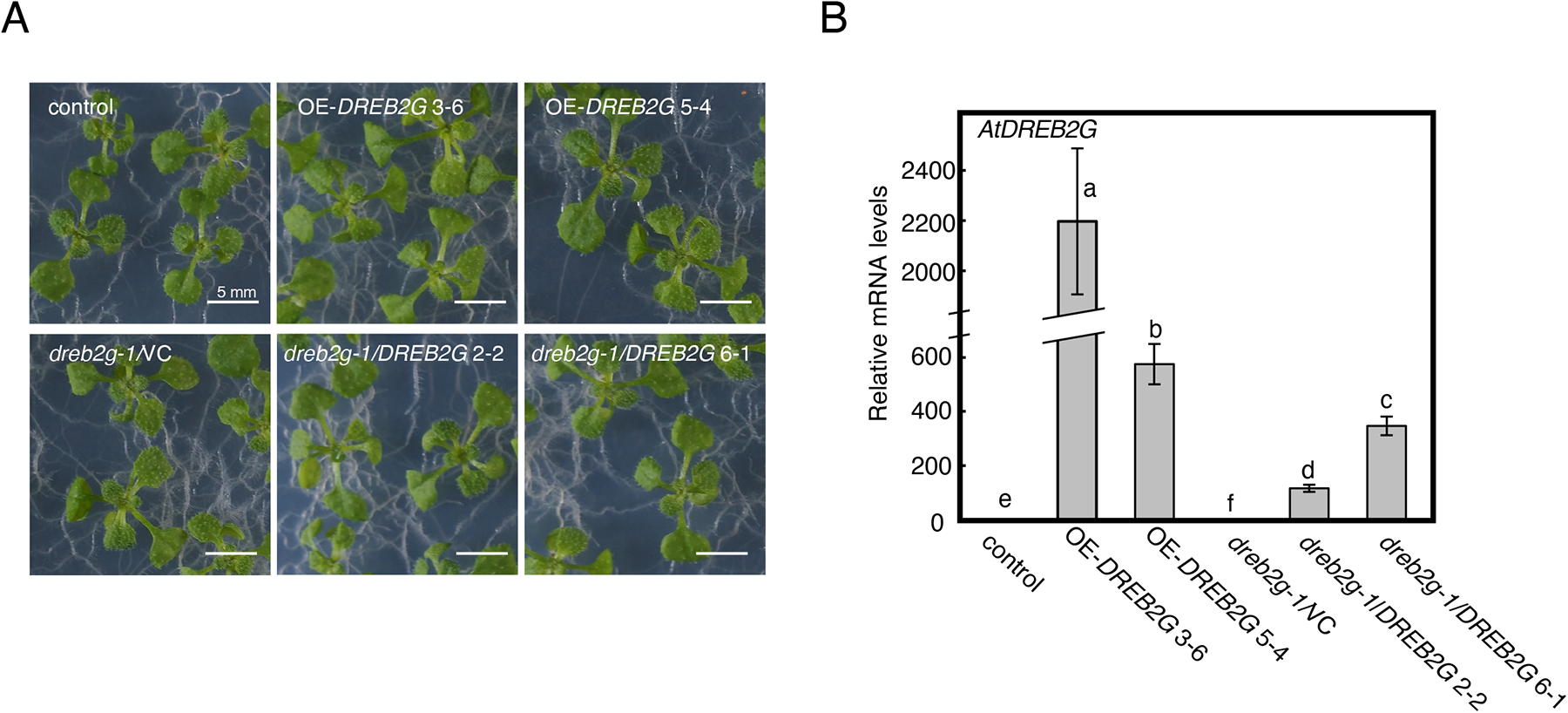
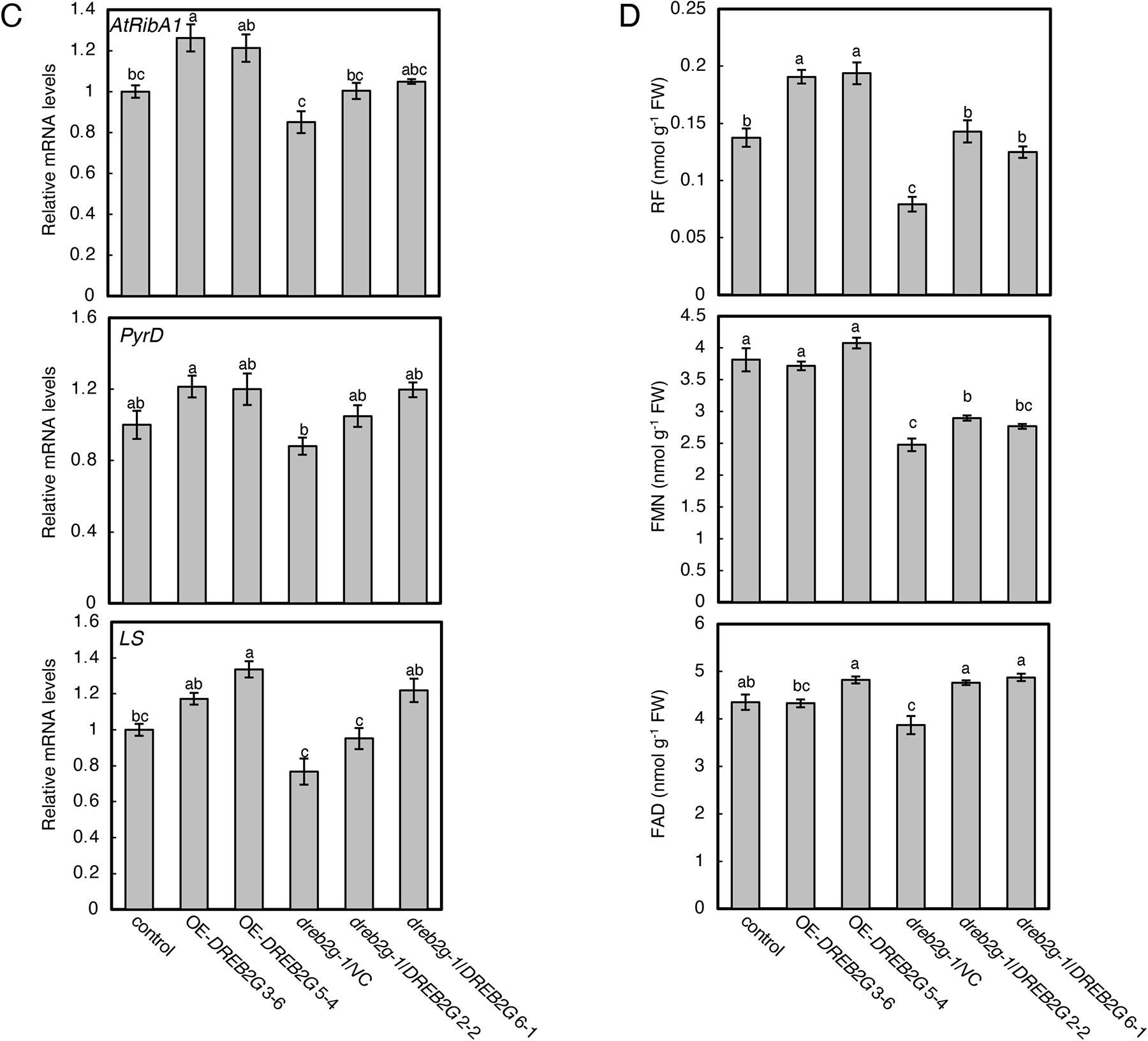
Characterization of the transgenic plants with constitutively overexpressing AtDREB2G in the wild-type and dreb2g-1 backgrounds. The control and *dreb2g-1*/VC (wild-type and *dreb2g-1* plants transformed with the empty vector, respectively), two independent lines overexpressing *AtDREB2G* in the wild-type background (OE-*DREB2G* 3-6 and 5-4), and two independent lines overexpressing *AtDREB2G* in the *dreb2g-1* background (*dreb2g-1*/*DREB2G* 2-2 and 6-1) were used. These plants were cultivated on 1/2 MS medium with 1% sucrose for two weeks under standard conditions. (A) Representative phenotype of transgenic lines. The transcript levels of (B) *AtDREB2G* and (C) RF biosynthetic genes. (D) Cellular flavin levels. The presented data represent means ± SE from a minimum of three individual experiments (n ≥ 3) involving independently grown plants. Different letters indicate significant differences (*P* < 0.05, Tukey-Kramer test).

To evaluate the influence of *AtDREB2G* on freezing tolerance, transgenic plants were subjected to -20°C and then returned to standard growth conditions. Notably, no significant differences in stress tolerance were observed among the control, *dreb2g-1*/VC, and *AtDREB2G*-overexpressing lines (Supplemental Fig. S5). Additionally, we assessed the ABA sensitivity of the transgenic lines. When seeds were germinated on 1/2 MS medium without ABA, there were no discernible differences in early development, as evidenced by germination rate and cotyledon greening rate, among the control, *dreb2g-1*/VC, OE-*DREB2G*, and *dreb2g-1*/*DREB2G* plants (Fig. 6A and 6B). In contrast, when seeds were germinated on a medium containing 0.5 μM ABA, the germination rate was significantly lower in *dreb2g-1*/VC plants, but higher in OE-*DREB2G* plants compared to control plants (Fig. 6A). The ABA sensitivity of *dreb2g-1*/VC plants was mitigated by the overexpression of *AtDREB2G* in the *dreb2g-1*/*DREB2G* plants. As judged by the cotyledon greening rate, OE-*DREB2G* seedlings displayed significant insensitivity to ABA treatment, while *dreb2g-1*/VC seedlings exhibited slightly enhanced sensitivity in comparison to control seedlings (Fig. 6B). These findings clearly illustrate the impact of *AtDREB2G* on ABA response in *Arabidopsis*.

**Fig. 6.**
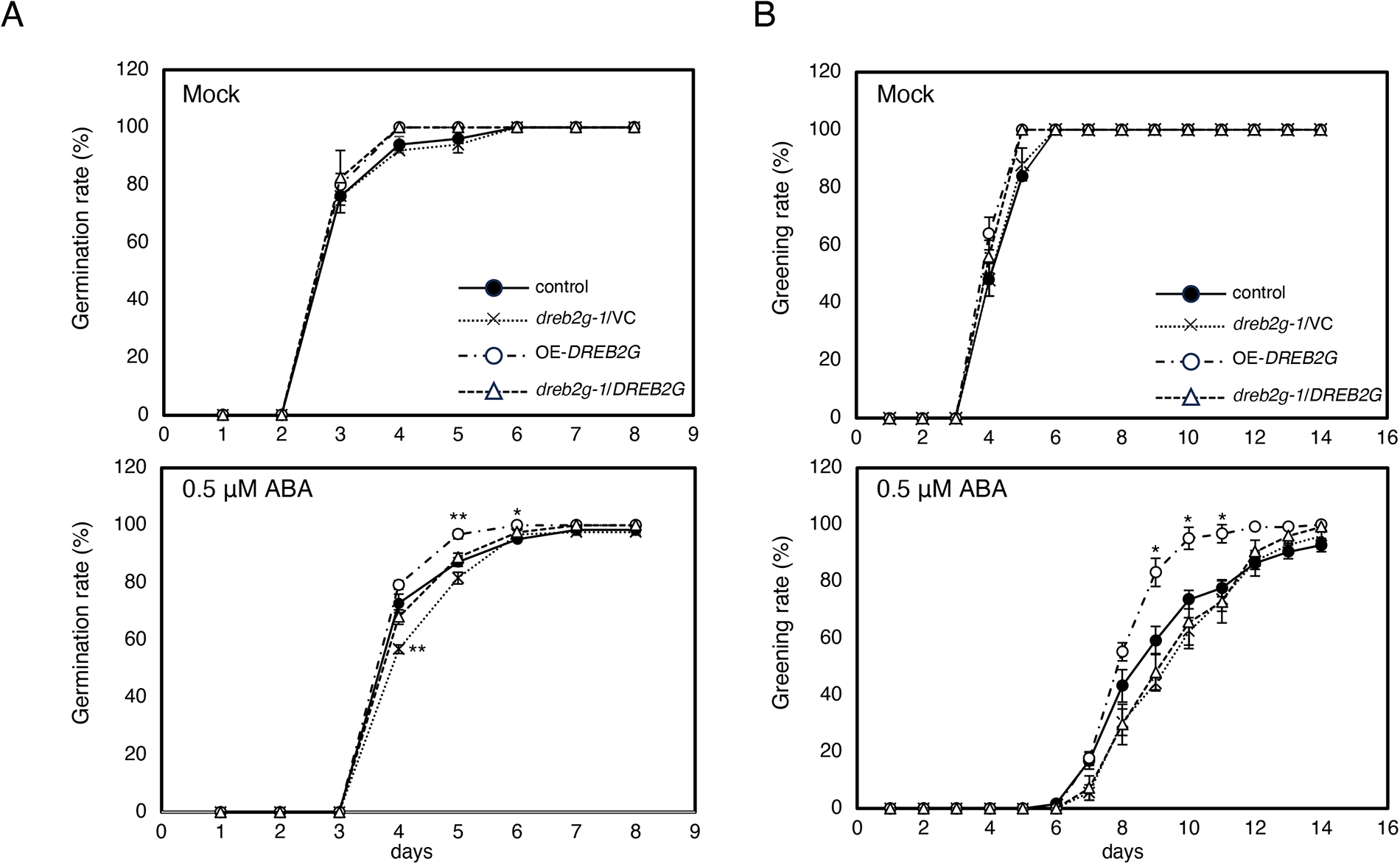
Sensitivity of *AtDREB2G* overexpressing lines to ABA treatment. Germination rate (A) and greening rate (B) of the control, *dreb2g-1*/VC, OE-*DREB2G* 3-6, and *dreb2g-1*/*DREB2G* 6-1 plants on 1/2 MS medium with or without 0.5 μM ABA. The germination rate represents the percentage of seeds in which the primary root has successfully penetrated the seed coat, while the greening rate indicates the percentage of seeds that have developed green cotyledons. The presented data represent means ± SE from a minimum of three individual experiments (n ≥ 3) involving independently grown plants. Asterisks indicate significant differences from the values of the wild-type plants (\**P* < 0.05 and \*\**P* < 0.01, Tukey-Kramer test).

## Discussion

In this study, we identified *AtDREB2G* as a novel regulator of flavin metabolism. Notably, *dreb2g-1* exhibited markedly reduced transcript levels of RF biosynthetic genes and lower cellular flavin levels compared to those in wild-type plants. These alterations were effectively complemented by the overexpression of *AtDREB2G*. Furthermore, both transient and constitutive overexpression of *AtDREB2G* resulted in elevated transcript levels of RF biosynthetic genes and increased cellular flavin levels. These findings unequivocally establish AtDREB2G as a positive regulator of RF biosynthesis in *Arabidopsis*. It is worth noting that our previous research suggested that flavin metabolism is subject to negative feedback regulation by cellular flavin levels (Maruta et al., 2012). However, it is unlikely that *AtDREB2G* participates in this process, as its expression was increased approximately twofold by treatment with 0.1 mM FAD compared to the mock treatment (Supplemental Table S1).

The transcription of *AtDREB2G* was significantly upregulated in response to low temperature and ABA treatment (Figs. 3 and 4). Concurrently, the transcription of genes involved in flavin biosynthesis and cellular flavin levels exhibited significant increases in wild-type plants under these conditions. Such increases were, however, compromised in *dreb2g-1* mutants (Figs. 3 and 4). These findings suggest that *AtDREB2G* plays a role in regulating flavin biosynthesis under low-temperature stress. Given the central role of ABA in cold acclimation, the induction of *AtDREB2G* transcription under low-temperature stress might be mediated through the ABA pathway, but the current study has not tested this possibility. The physiological significance of the *AtDREB2G*-mediated regulation of flavin metabolism under low-temperature stress remains unclear. Considering the essential role of flavin in various metabolic processes, including antioxidant networks, it might be plausible that the demand for flavin increases during low-temperature stress. Investigating the role of flavin metabolism in cold stress acclimation warrants future research. Interestingly, even though *AtDREB2G* transcription was induced under high-temperature stress, this stress led to reduced RF levels in both wild-type and *dreb2g-*1 plants (Supplemental Fig. S4). Several possibilities could explain this phenomenon, such as the inhibition of AtDREB2G’s function under high-temperature stress or the involvement of unknown regulators. Alternatively, it is conceivable that flavins may undergo degradation or instability under high-temperature stress conditions.

Further investigations are required to understand whether AtDREB2G directly regulates flavin metabolism. While the target genes of AtDREB1A and AtDREB2A have been identified using their overexpressing lines (Maruyama et al., 2004; Sakuma et al., 2006b), these do not include RF biosynthetic genes. Our search for DREs in the 1000-bp promoter regions of RF biosynthetic genes using PlantCARE, a database of plant *cis*-acting regulatory elements (Lescot et al., 2002), found that only one DRE in the promoter region of *PyrD*, with none found in *AtRibA1*, *PyrR*, *PyrP2*, *LS*, and *RS*. Assuming that AtDREB2G regulates the transcription of target genes by binding to DREs, it appears that the regulation of flavin biosynthesis by this transcription factor is indirect. Furthermore, we identified at least one ABA-responsive *cis*-acting element known as ABRE (ABA-responsive element) in the promoter regions of *PyrD*, *PyrP2*, *LS*, *RS*, and *AtDREB2G*. ABA indeed activated the transcription of these genes, suggesting the pivotal role of ABA in flavin metabolism. We observed that *dreb2g-1* mutants displayed high sensitivity to ABA treatment, which was alleviated by *AtDREB2G* overexpression in *dreb2g-1*/*DREB2G* plants (Figure 6). Conversely, OE-*DREB2G* plants exhibited insensitivity to ABA treatment (Figure 6). Similar results have been reported for transgenic *Arabidopsis* plants expressing *Lilium longiflorum DREB1G*, which also displayed reduced sensitivity to exogenous ABA treatment compared to wild-type plants (Liu et al., 2019). Hence, it is plausible that *AtDREB2G* negatively regulates ABA response. While it remains unclear whether the regulation of flavin metabolism by DREB is linked to ABA response, a maize mutant known as *defective kernel33* (*dek33*), causing abnormal seed development, suggests a potential relationship between flavin synthesis and ABA response. This mutant bears a mutation in the gene encoding pyrimidine reductase, a component of the RF biosynthetic pathway, resulting in decreased levels of both RF and ABA (Dai et al., 2019).

In conclusion, this study highlights the involvement of AtDREB2G in regulating flavin biosynthesis under conditions of low-temperature stress and ABA treatment. This provides new insights into the role of flavin biosynthesis regulation in responses to stress and hormones. Despite its fundamental importance, knowledge regarding flavin synthesis and function in plants remains notably limited. Several prior studies have suggested a connection between flavins and responses to stress and hormones. For instance, the *Arabidopsis* photosensitive mutant (*phs1*), lacking *PyrR*, exhibits low FAD levels, resulting in stunted growth, bleached leaves, and reduced ferredoxin-NADP^+^ oxidoreductase activity (a flavoenzyme) under high-light conditions (Ouyang et al., 2010). Additionally, a point mutation in the *COS1* (*coi1 suppressor1*) gene, encoding LS, suppresses the jasmonic acid-insensitive phenotype in *Arabidopsis coronatine insensitive1* (*coi1*) mutants (Xiao et al., 2004). As jasmonic acid is a stress hormone integral to responses to both abiotic and biotic stresses, *coi1 cos1* double mutants are highly susceptible to pathogens (Xiao et al., 2004). Moreover, external RF treatment activates the expression of *PR* genes, induces systemic acquired resistance, and enhances plant resistance to pathogens (Dong and Beer, 2000; Zhang et al., 2009). Expanding research into the function and regulation of flavin metabolism will be imperative for gaining a comprehensive understanding of plant responses to stress and hormones.

## Materials and methods

### Plant materials and growth conditions

*Arabidopsis thaliana* ecotype Col-0 was used as the wild-type plant. T-DNA insertion mutants (Supplemental Table S2) and a transgenic line in which *AtDREB2G* expression is estradiol-inducible (CS2102394) were obtained from the Arabidopsis Biological Resource Center. Seeds were sown in soil or on half-strength Murashige and Skoog (MS) medium containing 1% sucrose. After three days of incubation at 4°C in darkness, the seedlings were transferred to a growth chamber under standard conditions (16 hours of light at 22°C and 8 hours of darkness at 20°C), with a light intensity of 100 µmol photons m^-2^ s^-1^.

For freezing stress treatments, plants were initially grown on 1/2 MS medium agar plates for 10 days under standard growth conditions and were subsequently transferred to -20°C for 5 hours. Following stress exposure, the plants were returned to a growth chamber under standard conditions and allowed to grow for three days. In the case of exogenous FAD treatment, leaves from 3-week-old wild-type *Arabidopsis* plants cultivated in soil were detached and placed in dishes with distilled water, either with 0.1 mM FAD or without (mock), at 4 hours after the start of illumination. Subsequently, the leaves were gently rinsed three times with distilled water, promptly frozen in liquid nitrogen, and stored at -80°C until further analysis.

### Measurement of intracellular flavins

The frozen leaves of *Arabidopsis* plants (∼50 mg) were powdered using a multi-bead shocker (Yasui Kikai, Osaka, Japan) and then homogenized with 0.5 ml of 50% methanol. The resulting extracts were subjected to heating at 80°C for 10 minutes in darkness. After centrifugation at 20,000 × g and 4°C for 15 minutes, the supernatants were dried using a rotary evaporator in darkness and subsequently reconstituted in the mobile phase. These extracted samples were then filtered through a 0.2 μm filter (Millipore) and analyzed *via* HPLC, employing a COSMOSIL 5C18-MS-II column (4.6 x 250 mm, Nacalai Tesque, Kyoto, Japan) at a flow rate of 0.5 ml min-1, with a mobile phase consisting of 10 mM NaH_2_PO_4_ (pH 5.5) containing 30% methanol (v/v). To determine flavin contents in the extracts, fluorescence detection was employed with an excitation wavelength of 445 nm and an emission wavelength of 530 nm.

### Microarray analysis

Total RNA was extracted from the leaves following either mock or exogenous FAD treatment, using Sepasol-RNA I (Nacalai Tesque, Kyoto, Japan). The quality and purity of the RNA samples were verified for microarray analysis using an Ultrospec 2100 pro (GE Healthcare UK Ltd, Buckinghamshire, UK). Subsequently, the total RNA samples were reverse-transcribed to generate double-stranded cDNA, which was further transcribed *in vitro* in the presence of biotin-labeled nucleotides using an IVT Labeling Kit (Affymetrix Inc.). The labeled cRNA was then fragmented and subjected to hybridization on Affymetrix ATH1 GeneChip arrays at 45°C for 16 hours, following Affymetrix protocols. Arrays were subjected to washing on the Affymetrix Fluidics Station 450 and fluorescence intensity measurements were obtained using an Affymetrix GeneChip Scanner 3000. Raw data were processed using Affymetrix Gene Chip Operating Software (GCOS; v1.4.0.036). Two biological replicates were included in the analysis. The microarray data have been deposited in the public NCBI Gene Expression Omnibus database under the GEO accession number GSE169061.

### Semi-quantitative and quantitative real-time PCR analysis

Total RNA extracted from *Arabidopsis* leaves (see above) was reverse transcribed into first-strand cDNA using ReverTra Ace (Toyobo) with the oligo (dT) 20 primer. In the semi-quantitative RT-PCR analysis, PCR amplification was conducted through 22 to 35 cycles of denaturation at 95°C for 30 seconds, annealing at 55°C for 30 seconds, and extension at 72°C for 60 seconds, followed by a final extension step at 72°C for 10 minutes. The uniform loading of each amplified cDNA was verified using the Actin8 control PCR product. Quantitative real-time PCR analysis was carried out following the procedures outlined in a previous study (Ogawa et al. 2016). Primer sequences are provided in Supplemental Tables S2 and S3.

### ES treatment

Transgenic seeds obtained from ABRC (CS2102394) were planted on half-strength MS medium containing 1% sucrose. The plates were subjected to a 3-day period of stratification in the dark at 4°C, followed by transfer to a growth chamber under standard growth conditions. After 2 weeks, the seedlings were moved to 1/2 MS medium containing 2 μM ES. Shoots were harvested and utilized for q-PCR and flavin content analyses.

### Plasmid construction and transformation of Arabidopsis

To construct the plasmid for the constitutive overexpression of *AtDREB2G*, we employed a modified pRI201-AN vector (Takara, Kyoto, Japan) in which the *NPTII* gene was substituted with the *HPT* gene. AtDREB2G cDNA was amplified from the first-strand cDNA using the following primer sets: pRI201-At5g18450-F (5’-CTGTTGATACATCACATGGAAGAAGAGCAACCTCC-3’) and pRI201-At5g18450-R (5’-TGTCGAGGTCGACACTCAGAACCAATTCCATGGAT-3’). The amplified cDNA was inserted downstream of the CaMV 35S promoter in the modified pRI201-AN vector using In-Fusion Cloning technology (Takara). *Agrobacterium tumefaciens* strain C58, transformed with the construct obtained through electroporation, was used to infect *Arabidopsis* wild-type plants (Col-0) and *dreb2g-1* mutants using the floral dip transformation method. T_1_ seedlings were chosen on half-strength MS medium in Petri dishes supplemented with 1% sucrose and 20 mg L^−1^ hygromycin for 2 weeks before being transferred to soil.

### ABA sensitivity analysis

Seeds were planted on 1/2 MS medium containing 1% sucrose and 0.5 μM ABA and kept in darkness at 4 °C for three days. Subsequently, the plants were cultivated in a growth chamber under standard conditions. Germination and cotyledon greening rates were recorded at various time points. Germination was considered when primary roots emerged through the seed coat, while the greening rate represented the proportion of total seeds that developed green cotyledons.

### Statistical analyses

The statistical analyses of the data were based on Dunnett’s, Tukey-Kramer, or Student’s *t*-test (see figure legends). All calculations were performed using at least three independent biological replicates.

## Abbreviations

FAD: flavin adenine dinucleotide
FMN: flavin mononucleotide
RF: riboflavin
ABA: abscisic acid
DREB: dehydration responsive element binding protein

## Data Availability

The data underlying this work are available in the article and supplementary materials.

## Author Contributions

K.Y., S.S., and T.O. conceived the research plans. J.N. and M.H. performed most of the experiments. Y.T. performed microarray analysis. T.M., T.I., and T.O. supervised the experiments. J.N., M.H., and T.O. designed the experiments. J.N., M.H., and T.O. analyzed the data. K.Y. and T.O. wrote the article. All authors have read and agreed to the published version of the manuscript.

