## Supplemental Figs for "AtDREB2G is a novel regulator of riboflavin biosynthesis under low-temperature stress and abscisic acid treatment in *Arabidopsis thaliana*"

### Slide 1
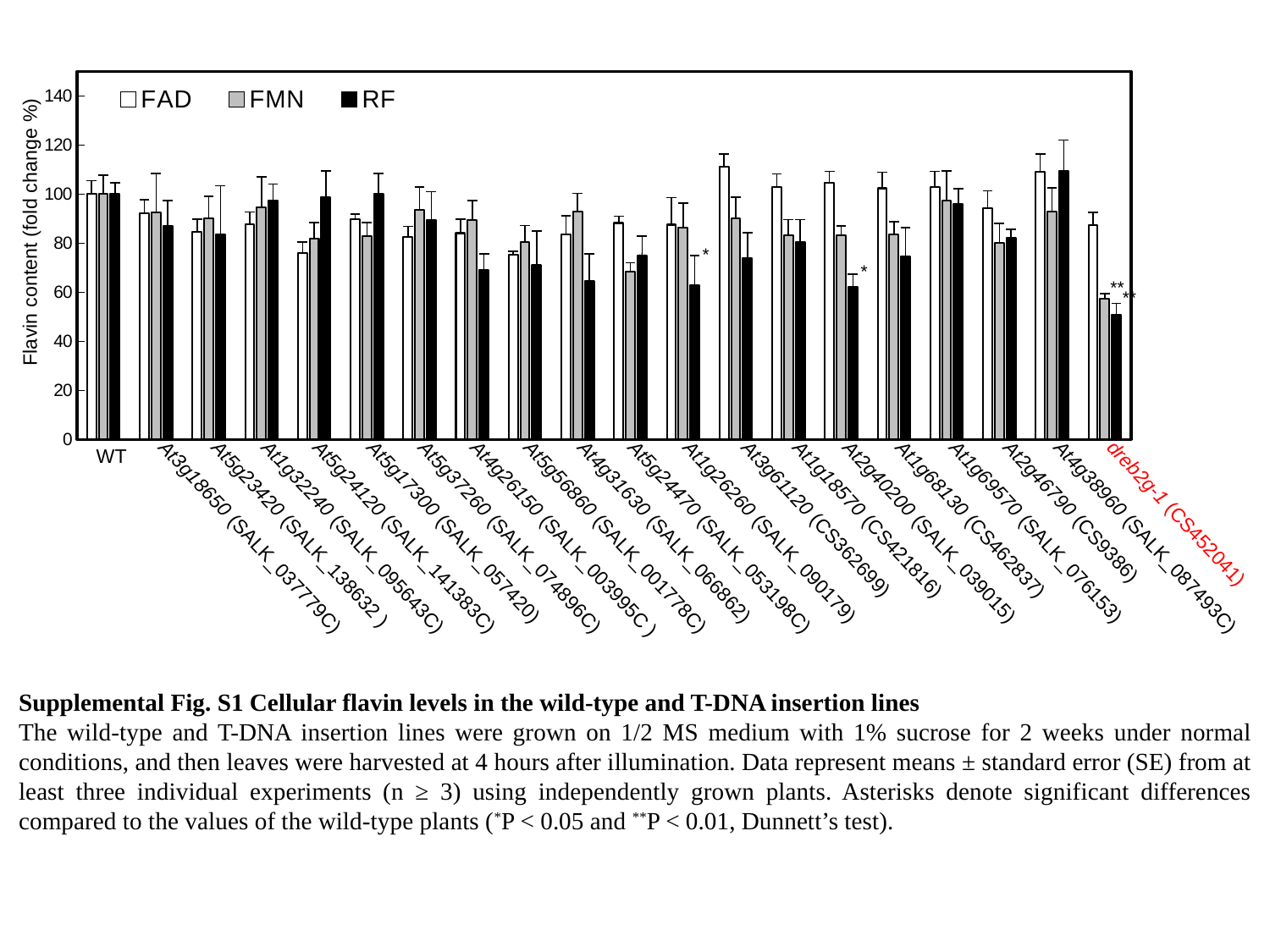

#### Chart
| Category | | | |
|---|---|---|---|
| WT | 100.0 | 100.0 | 100.0 |
| frtf1 | 92.2290439094016 | 92.61028584527773 | 87.05842844826482 |
| frtf2 | 84.59806322692486 | 90.13476995689987 | 83.44385159175917 |
| frtf3 | 87.78767334252335 | 94.51863051016701 | 97.43453891219066 |
| frtf4 | 75.94873493800803 | 81.71282382251395 | 98.67268143774545 |
| frtf5 | 89.7782995381715 | 82.919530854875 | 100.0565513815839 |
| frtf6 | 82.3501740196297 | 93.6283706593363 | 89.30844520475394 |
| frtf7 | 84.06753677694836 | 89.28579104081143 | 68.9117838048438 |
| frtf8 | 75.20025795896359 | 80.28954487329909 | 70.90421203386356 |
| frtf9 | 83.55547112682035 | 92.71191606157925 | 64.46929200921258 |
| frtf10 | 88.19294160608808 | 68.48856696510485 | 75.01298054660158 |
| frtf11 | 87.50566854139416 | 86.43276002572219 | 62.84106213097827 |
| frtf12 | 111.20000305891664 | 90.02017727124222 | 74.04147615314132 |
| frtf13 | 102.89156536564839 | 83.2871182813293 | 80.34139665970926 |
| frtf14 | 104.65300585431513 | 83.11646967296139 | 62.00018278300231 |
| frtf15 | 102.34446767210042 | 83.39616871217007 | 74.60150378015811 |
| frtf16 | 102.97768041375933 | 97.17400515217507 | 96.04362816146413 |
| frtf17 | 94.34144994415675 | 80.19399172480091 | 82.0937966437853 |
| frtf18 | 109.21453943920044 | 92.73990985411344 | 109.34399466527313 |
| frtf19.1 | 87.4460184501604 | 57.23609115223869 | 50.72771320446846 |Flavin content (fold change %)
*
*
*
**
**
WT
At2g46790 (CS9386)
dreb2g-1 (CS452041)
At3g61120 (CS362699)
At1g18570 (CS421816)
At1g68130 (CS462837)
At5g17300 (SALK_057420)
At4g31630 (SALK_066862)
At1g26260 (SALK_090179)
At2g40200 (SALK_039015)
At1g69570 (SALK_076153)
At5g23420 (SALK_138632 )
At3g18650 (SALK_037779C)
At1g32240 (SALK_095643C)
At5g24120 (SALK_141383C)
At5g37260 (SALK_074896C)
At5g56860 (SALK_001778C)
At5g24470 (SALK_053198C)
At4g38960 (SALK_087493C)
At4g26150 (SALK_003995C )
Supplemental Fig. S1 Cellular flavin levels in the wild-type and T-DNA insertion lines
The wild-type and T-DNA insertion lines were grown on 1/2 MS medium with 1% sucrose for 2 weeks under normal conditions, and then leaves were harvested at 4 hours after illumination. Data represent means ± standard error (SE) from at least three individual experiments (n ≥ 3) using independently grown plants. Asterisks denote significant differences compared to the values of the wild-type plants (^*^P < 0.05 and ^**^P < 0.01, Dunnett’s test).

### Slide 2
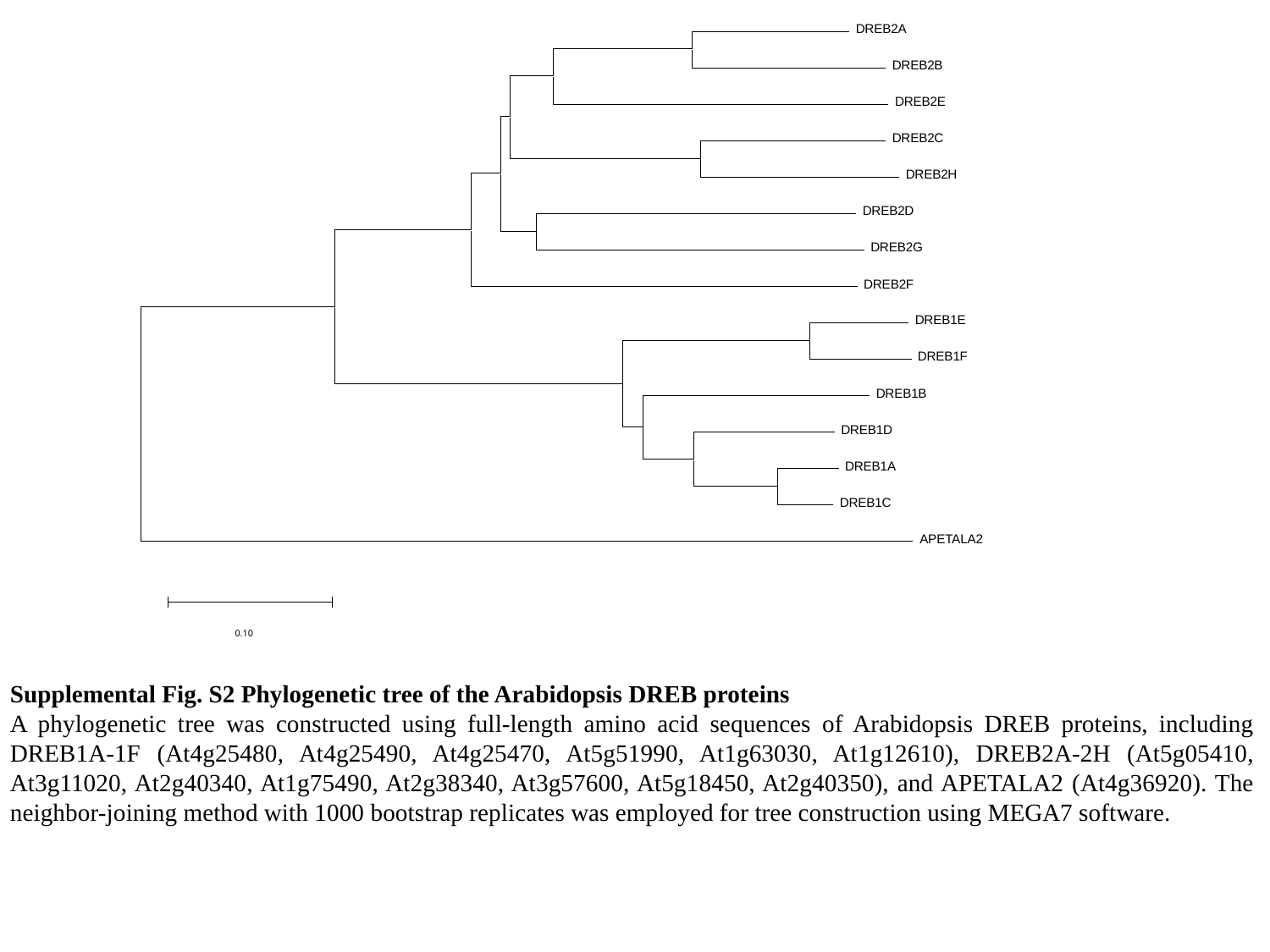

DREB2A
 DREB2B
 DREB2E
 DREB2C
 DREB2H
 DREB2D
 DREB2G
 DREB2F
 DREB1E
 DREB1F
 DREB1B
 DREB1D
 DREB1A
 DREB1C
 APETALA2
0.10
Supplemental Fig. S2 Phylogenetic tree of the Arabidopsis DREB proteins
A phylogenetic tree was constructed using full-length amino acid sequences of Arabidopsis DREB proteins, including DREB1A-1F (At4g25480, At4g25490, At4g25470, At5g51990, At1g63030, At1g12610), DREB2A-2H (At5g05410, At3g11020, At2g40340, At1g75490, At2g38340, At3g57600, At5g18450, At2g40350), and APETALA2 (At4g36920). The neighbor-joining method with 1000 bootstrap replicates was employed for tree construction using MEGA7 software.

### Slide 3
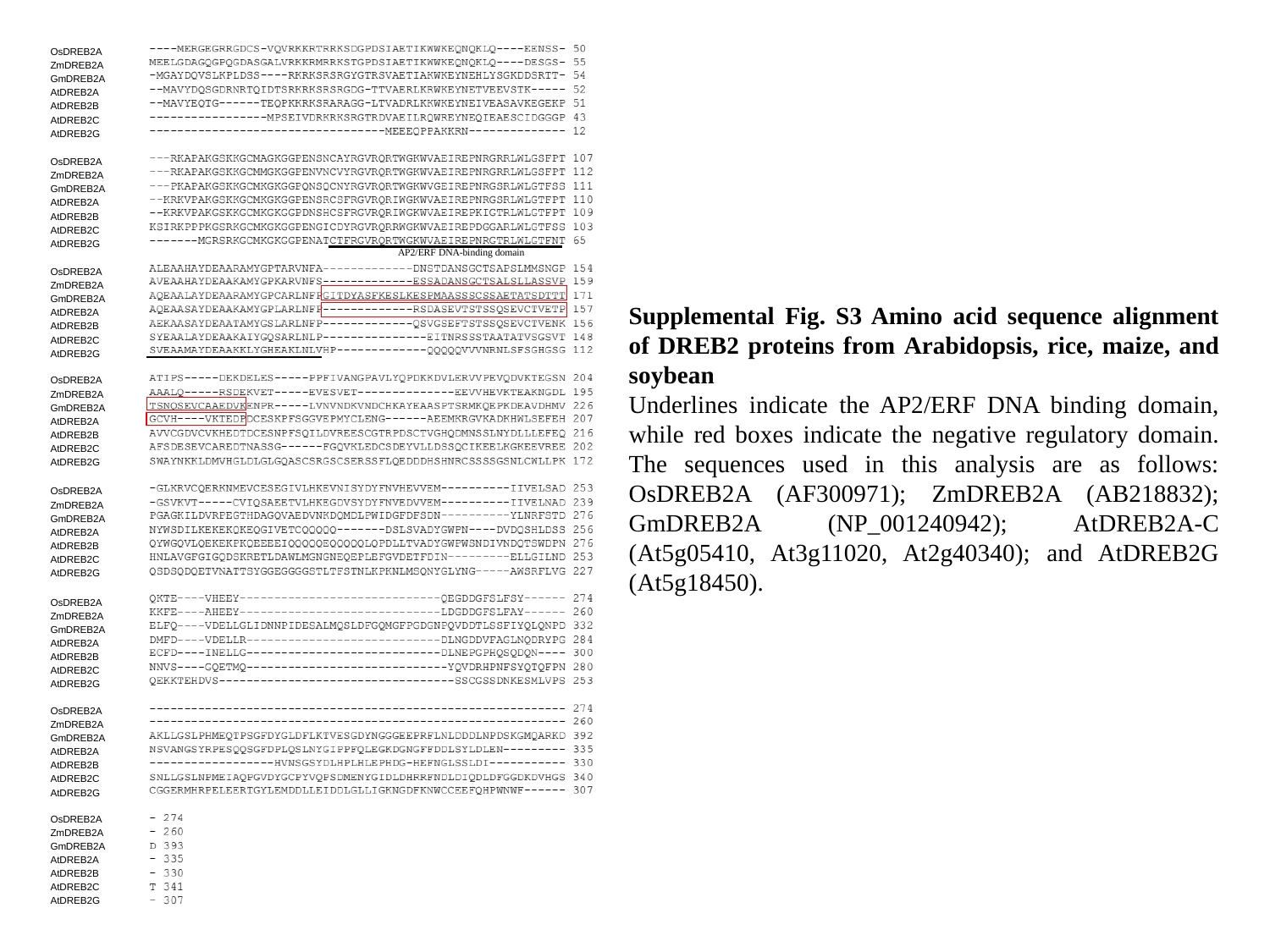

OsDREB2A
ZmDREB2A
GmDREB2A
AtDREB2A
AtDREB2B
AtDREB2C
AtDREB2G
OsDREB2A
ZmDREB2A
GmDREB2A
AtDREB2A
AtDREB2B
AtDREB2C
AtDREB2G
AP2/ERF DNA-binding domain
OsDREB2A
ZmDREB2A
GmDREB2A
AtDREB2A
AtDREB2B
AtDREB2C
AtDREB2G
Supplemental Fig. S3 Amino acid sequence alignment of DREB2 proteins from Arabidopsis, rice, maize, and soybean
Underlines indicate the AP2/ERF DNA binding domain, while red boxes indicate the negative regulatory domain. The sequences used in this analysis are as follows: OsDREB2A (AF300971); ZmDREB2A (AB218832); GmDREB2A (NP_001240942); AtDREB2A-C (At5g05410, At3g11020, At2g40340); and AtDREB2G (At5g18450).
OsDREB2A
ZmDREB2A
GmDREB2A
AtDREB2A
AtDREB2B
AtDREB2C
AtDREB2G
OsDREB2A
ZmDREB2A
GmDREB2A
AtDREB2A
AtDREB2B
AtDREB2C
AtDREB2G
OsDREB2A
ZmDREB2A
GmDREB2A
AtDREB2A
AtDREB2B
AtDREB2C
AtDREB2G
OsDREB2A
ZmDREB2A
GmDREB2A
AtDREB2A
AtDREB2B
AtDREB2C
AtDREB2G
OsDREB2A
ZmDREB2A
GmDREB2A
AtDREB2A
AtDREB2B
AtDREB2C
AtDREB2G

### Slide 4
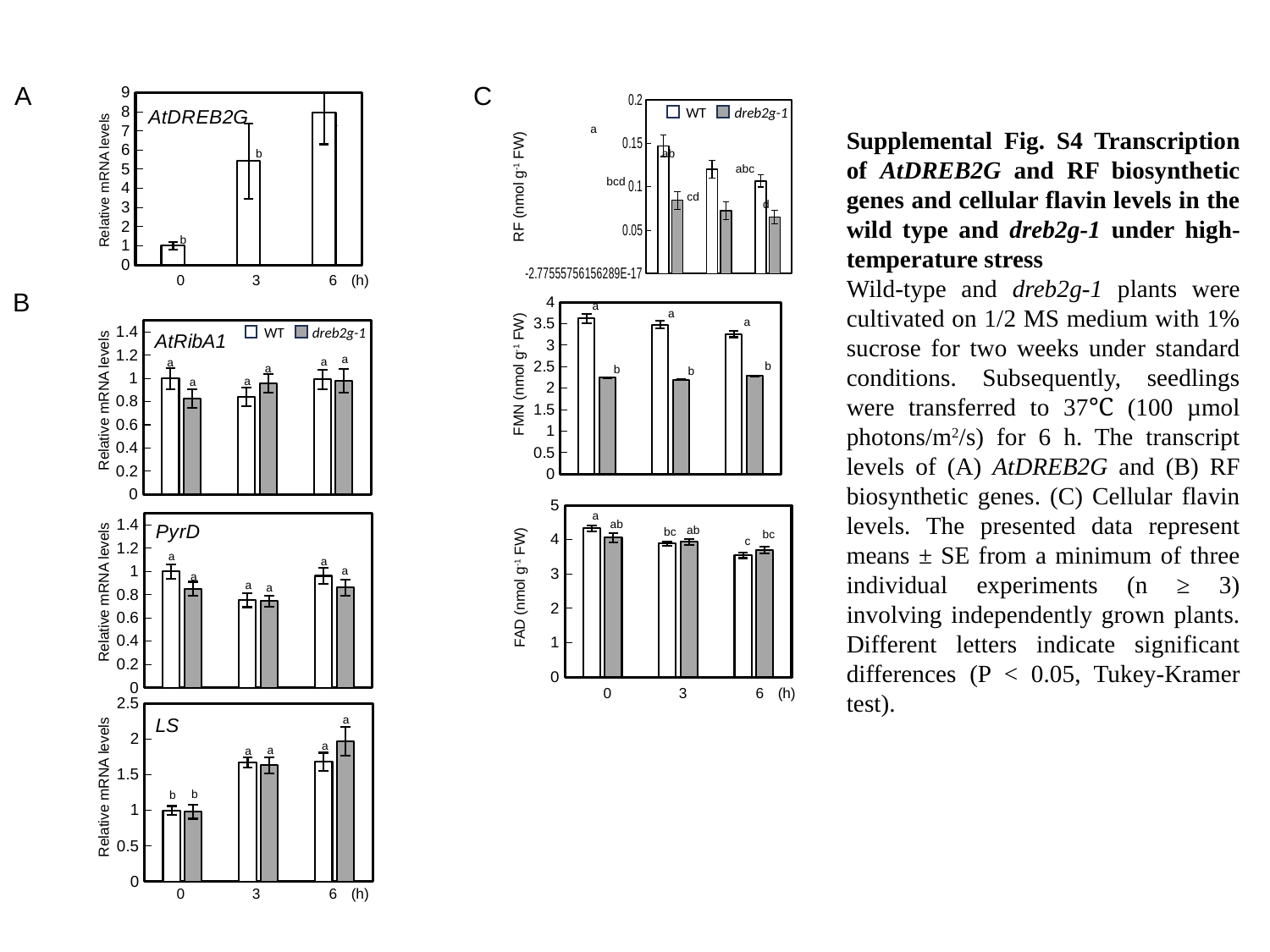

A
C
#### Chart: AtDREB2G
| Category | |
|---|---|
| 0h | 1.0 |
| 3h | 5.427029586616043 |
| 6h | 7.94148060367181 |
#### Chart
| Category | | |
|---|---|---|
| 0h | 0.14699753465614768 | 0.08441285638817765 |
| 3h | 0.12006069946285575 | 0.07241906856119924 |
| 6h | 0.1065739051508718 | 0.06526733370962919 |WT
dreb2g-1
a
a
Supplemental Fig. S4 Transcription of AtDREB2G and RF biosynthetic genes and cellular flavin levels in the wild type and dreb2g-1 under high-temperature stress
Wild-type and dreb2g-1 plants were cultivated on 1/2 MS medium with 1% sucrose for two weeks under standard conditions. Subsequently, seedlings were transferred to 37℃ (100 µmol photons/m^2^/s) for 6 h. The transcript levels of (A) AtDREB2G and (B) RF biosynthetic genes. (C) Cellular flavin levels. The presented data represent means ± SE from a minimum of three individual experiments (n ≥ 3) involving independently grown plants. Different letters indicate significant differences (P < 0.05, Tukey-Kramer test).
b
ab
abc
Relative mRNA levels
bcd
RF (nmol g^-1^ FW)
cd
d
b
0
3
6
(h)
B
a
#### Chart
| Category | | |
|---|---|---|
| 0h | 3.6233347707266894 | 2.249495313359849 |
| 3h | 3.4782339150182553 | 2.1962130616746234 |
| 6h | 3.262586399440873 | 2.2816000802797025 |a
a
#### Chart: AtRibA1
| Category | | |
|---|---|---|
| 0h | 1.0 | 0.8270608245693887 |
| 3h | 0.8392209472246845 | 0.9552739381453179 |
| 6h | 0.9923364396425751 | 0.9789952806458568 |WT
dreb2g-1
a
a
a
b
a
b
b
FMN (nmol g^-1^ FW)
a
a
Relative mRNA levels
#### Chart
| Category | | |
|---|---|---|
| 0h | 4.330852149318878 | 4.0612661430158825 |
| 3h | 3.884120855479664 | 3.9424115007498406 |
| 6h | 3.547488672909259 | 3.7066243906088374 |
#### Chart: PyrD
| Category | | |
|---|---|---|
| 0h | 1.0 | 0.8500974219889751 |
| 3h | 0.7541000460184268 | 0.7458040006657985 |
| 6h | 0.962401966063858 | 0.8615236994898808 |a
ab
ab
bc
bc
c
a
a
a
a
a
FAD (nmol g^-1^ FW)
a
Relative mRNA levels
0
3
6
(h)
#### Chart: LS
| Category | | |
|---|---|---|
| 0h | 1.0 | 0.9816577023858777 |
| 3h | 1.6764308371259133 | 1.6330850917114879 |
| 6h | 1.682526548062336 | 1.9696593573300232 |a
a
a
a
Relative mRNA levels
b
b
0
3
6
(h)

### Slide 5
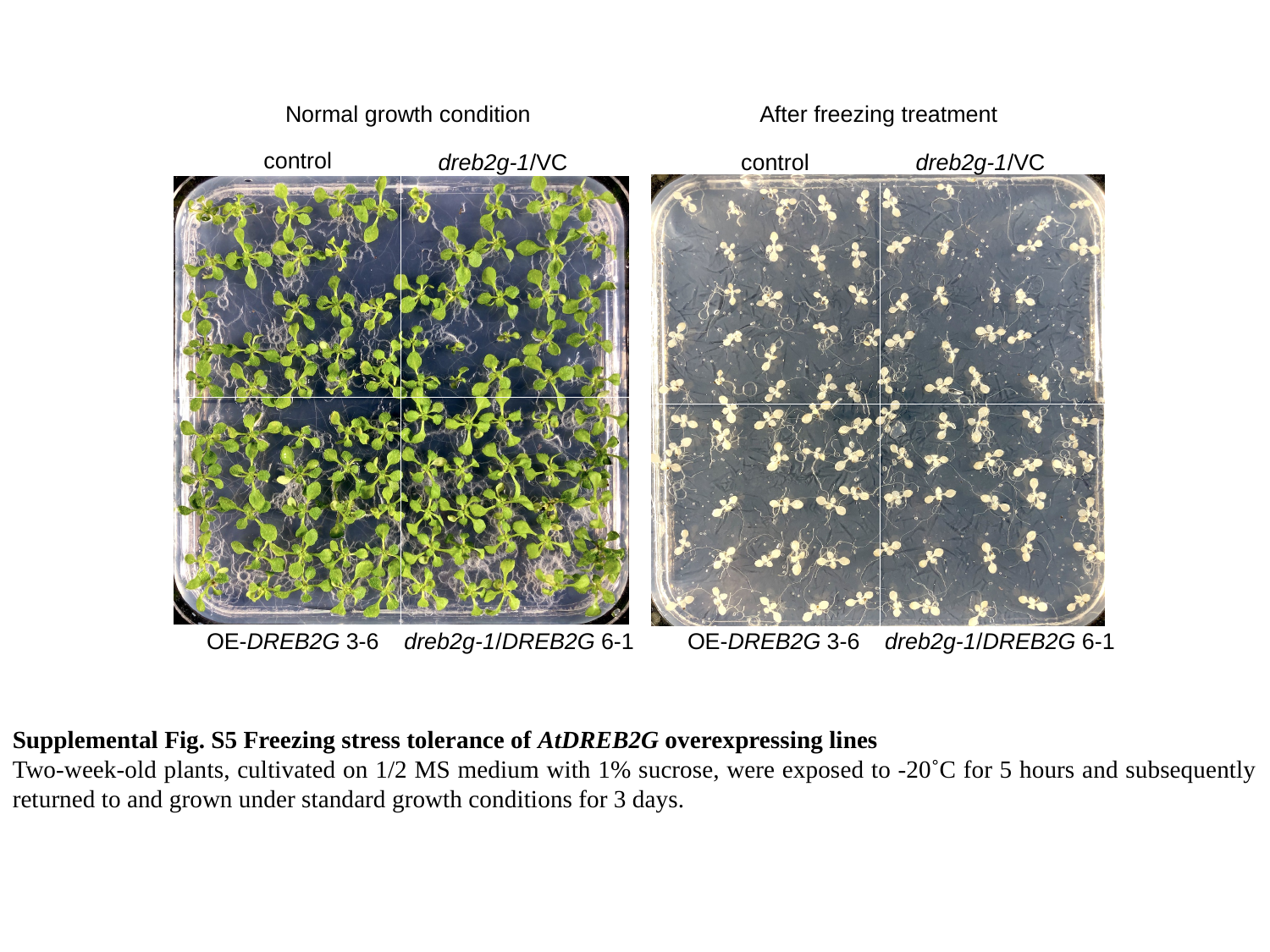

Normal growth condition
After freezing treatment
control
dreb2g-1/VC
control
dreb2g-1/VC
OE-DREB2G 3-6
dreb2g-1/DREB2G 6-1
OE-DREB2G 3-6
dreb2g-1/DREB2G 6-1
Supplemental Fig. S5 Freezing stress tolerance of AtDREB2G overexpressing lines
Two-week-old plants, cultivated on 1/2 MS medium with 1% sucrose, were exposed to -20˚C for 5 hours and subsequently returned to and grown under standard growth conditions for 3 days.
